# Low-frequency ERK and Akt activity dynamics are predictive of stochastic cell division events

**DOI:** 10.1101/2024.01.17.576041

**Authors:** Jamie J. R. Bennett, Alan D. Stern, Xiang Zhang, Marc R. Birtwistle, Gaurav Pandey

## Abstract

Understanding the dynamics of intracellular signaling pathways, such as ERK1/2 (ERK) and Akt1/2 (Akt), in the context of cell fate decisions is important for advancing our knowledge of cellular processes and diseases, particularly cancer. While previous studies have established associations between ERK and Akt activities and proliferative cell fate, the heterogeneity of single-cell responses adds complexity to this understanding. This study employed a data-driven approach to address this challenge, developing machine learning models trained on a dataset of growth factor-induced ERK and Akt activity time courses in single cells, to predict cell division events. The most effective predictive models were developed by applying discrete wavelet transforms (DWTs) to extract low-frequency features from the time courses, followed by using Ensemble Integration, an effective data integration and predictive modeling framework. The results demonstrated that these models effectively predicted cell division events in MCF10A cells (F-measure=0.524, AUC=0.726). ERK dynamics were found to be more predictive than Akt, but the combination of both measurements further enhanced predictive performance. The ERK model’s performance also generalized to predicting division events in RPE cells, indicating the potential applicability of these models and our data-driven methodology for predicting cell division across different biological contexts. Interpretation of these models suggested that ERK dynamics throughout the cell cycle, rather than immediately after growth factor stimulation, were associated with the likelihood of cell division. Overall, this work contributes insights into the predictive power of intra-cellular signaling dynamics for cell fate decisions, and highlights the potential of machine learning approaches in unraveling complex cellular behaviors.

## Introduction

Mammalian ERK1/2 (ERK) and Akt1/2 (Akt) kinases are ubiquitous regulators of proliferation, growth and survival [1, 2]. Their activity is typically controlled by upstream growth factor receptors (or other pathways affecting receptors [3, 4]), and deregulated by a variety of oncogenic (e.g. EGFR, HER2, RAS, BRAF and PI3K) and tumor suppressor (e.g. PTEN and NF1) mutations [5, 6, 7, 8, 9, 10, 11, 12, 13]. Thus, it is not surprising that these pathways are also important drug targets for treating cancers and other diseases [14, 15, 16, 17, 18, 19].

While prior work (e.g., reviewed by [1, 2]) established associations between ERK and Akt activity and proliferative cell fates, single-cell signaling and response heterogeneity makes this understanding more opaque. For example, it is known that clonal cells exposed to the same conditions can exhibit markedly different ERK and Akt activity dynamics and magnitudes, as well as different proliferative (and other) fates [20, 21, 22, 23, 24, 25, 26, 27, 28, 29, 30]. It has been hypothesized that temporal differences in kinase activity profiles, such as transient vs. sustained signaling [31, 32, 26, 33], oscillatory-like pulsing [23, 25, 24, 34, 22, 35, 36] and time-integrals [37, 20, 38, 27, 39] are associated with cell fate heterogeneity. Such kinetic control of cell fate decisions is not limited to the ERK and Akt pathways, but is also thought to be operative in systems such as p53 and NF*κ*B [40, 41, 42, 43, 44, 45, 46]. However, a question that remains open is whether any such dynamic features of ERK and Akt activity are predictive of proliferative fates in individual cells. Answering this question can be facilitated by experiments that track kinase activities in multiple single live cells with a matched readout of the cell fate, along with rigorous data science-based modeling that can robustly assess the links between the two.

Our previous study reported a dataset that coupled growth factor-induced ERK and Akt activity time courses to division events in the same single cells [27]. However, the question of predicting if a single, individual cell will divide or not based on such growth factor-induced time courses remains unexplored. Moreover, the previous study only considered a single cell line (MCF10A), a single set of growth factors (EGF and insulin), and acute, synchronized response to growth factors (as opposed to chronic, asynchronous scenarios). In the current work, we analyzed the previously generated dataset by utilizing time series transformations to extract features [47, 48, 49], and predicting cell divisions using effective machine learning techniques [50, 51, 52] for both MCF10A cell data with acute response to growth factors from a serum-starved state, and retinal pigment epithelial (RPE) cells asynchronously cycling in full growth media [53]. Models developed from only MCF10A data performed well on both datasets, and analyses suggested operational relationships between ERK and Akt dynamics and the probability of cell division.

## Results

### Data preparation

We first prepared our previously collected dataset [27] for the application of machine learning (ML) algorithms (Fig. 1A). Briefly, the dataset was generated from MCF10A cells expressing both ERK and Akt activity reporters [54, 55]. Serum and growth factor-starved cells were treated with EGF and insulin (two key components of MCF10A growth media), and imaged periodically for two days (Fig. 1A). For each cell, image analysis-extracted kinase activity time course data were collected along with cell fate, the latter enabling individual cells to be assigned the label of “divided” or “undivided” that we predicted via supervised ML methods [56] (Fig. 1A). We produced two sets of data from a “high-dose” and a “low-dose” experiment, where a higher/lower dosage of EGF and insulin were used, respectively. To enable rigorous model training and evaluation, the high-dose data were randomly divided in an 80:20 ratio into train and test sets, and the “low-dose” data were used as another test set. Table 1 provides a summary of these training and test sets.

**Table 1:** Summary of the datasets used in our study. Treatment was either epidermal growth factor (EGF) + Insulin in the case of serum-starved cells, or ERK inhibitor (ERKi) in the case of cells asynchronously cycling in full growth medium. Reporters were either kinase translocation reporters (KTR)-, or fluorescence resonance energy transfer (FRET)-based.

| Name | Treatment | Serum-starved? | Reporter | Number of cells |  |  | Cell type | Kinase(s) (modalities) |
| --- | --- | --- | --- | --- | --- | --- | --- | --- |
|  |  |  |  | Divided | Undivided | Total |  |  |
| High-dose (train) | 20 ng/mL EGF + 10 µg/mL Insulin | Yes | KTR | 246 | 756 | 1002 | MCF10A | ERK, Akt |
| High-dose (test) | 20 ng/mL EGF + 10 µg/mL Insulin | Yes | KTR | 62 | 189 | 251 | MCF10A | ERK, Akt |
| Low-dose (test) | 2 ng/mL EGF + 1 µg/mL Insulin | Yes | KTR | 246 | 1148 | 1394 | MCF10A | ERK, Akt |
| RPE (test) | 62.5 nM ERKi | No | FRET | 123 | 102 | 225 | RPE | ERK |

**Figure 1:**
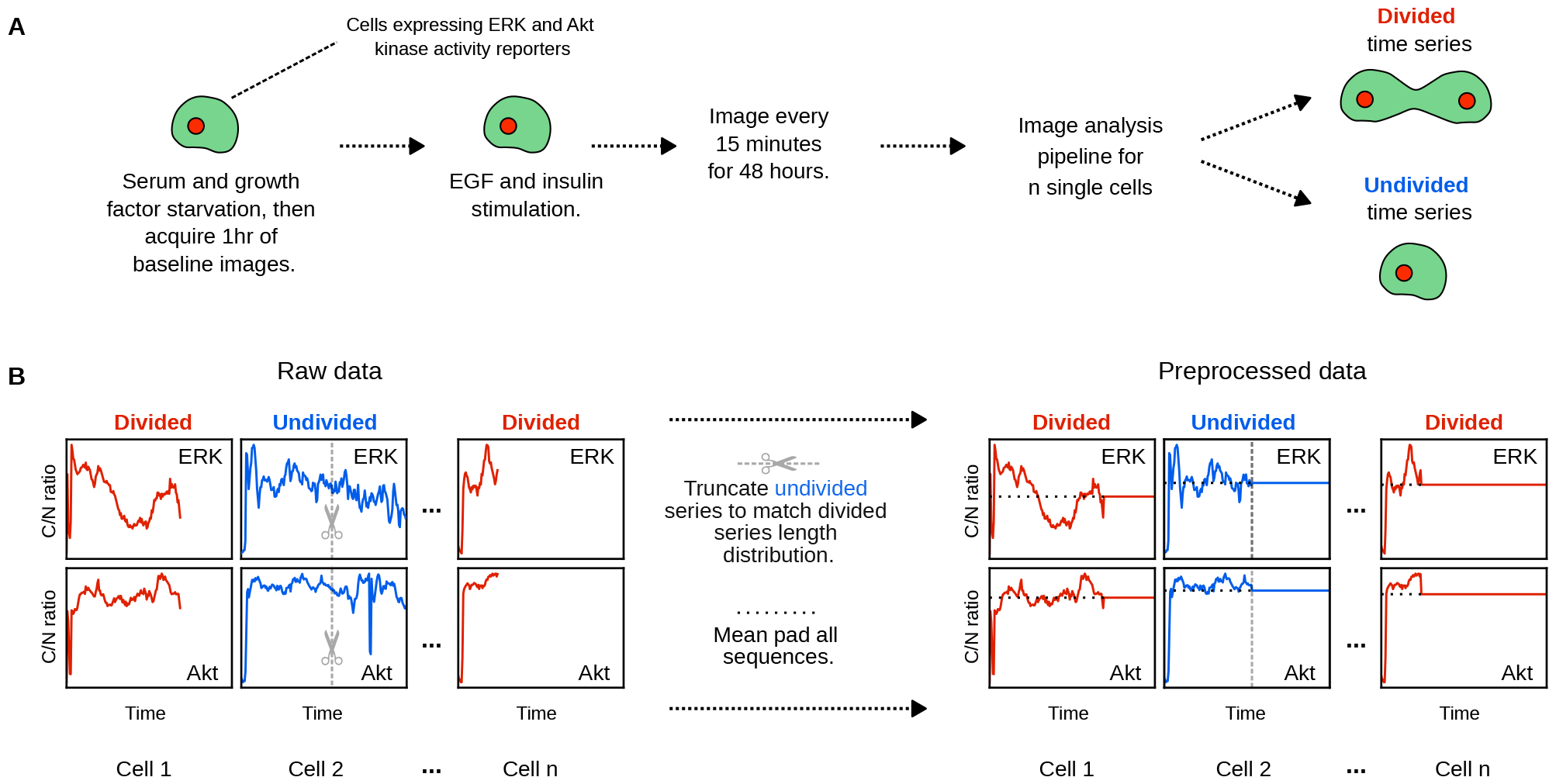
Overview of input data and their processing. **A**. Experimental workflow of MCF10A data generation: Initially, cells expressing ERK and Akt reporters were serum- and growth factor-starved. An hour of baseline images were taken immediately before growth factor treatment. After treatment with epidermal growth factor (EGF) and insulin, cells were imaged every 15 minutes for 2 days, and the resultant image time courses were analyzed to measure kinase activity (cytoplasmic to nuclear fluorescence ratio of the kinase translocation reporter (KTR) (C/N ratio)). Time courses were labeled as divided or undivided according to the fate of the corresponding cell. **B**. Pre-processing kinase time course data for machine learning analyses: Divided cells had a truncated time course at the time of cell division, whereas undivided cells did not. To address this incompatibility, all undivided cell time courses were truncated so that the distribution of time series lengths were the same between the two classes. Each time course was then padded to lengths equal to those of the undivided cell with the corresponding mean.

An important issue in these datasets was that divided cells had shorter time courses than undivided cells, since the original measurements terminated at the time of cell division [27]. This difference in time course length could have caused the downstream cell fate classifiers to trivially predict the outcome based on this artifact. Therefore, we processed the undivided cell time courses such that they were not trivially distinguishable from divided cell time courses (Fig. 1B; details in Methods and Supplementary Material).

### Discrete wavelet transforms combined with a heterogeneous ensemble yielded the most effective classifier of MCF10A cell division fate

We employed a multi-modal data fusion approach to leverage the likely complementary information contained within ERK and Akt time courses in order to predict cell fates. We compared a diverse range of integrative methods for predicting cell fate, and also compared them to predictions from the individual ERK and Akt time courses. Throughout this paper, we refer to multi-modal methods as [ERK, Akt], and single modality time courses by the name of the corresponding kinase.

Extracting structured features that capture important information in time courses is often an effective method to enable classification [57]. We evaluated several established transformation-based feature extraction techniques: discrete wavelet transforms (DWTs) [58], MiniRocket [59], tsfresh [60] and the amplitude of the Fourier transform (FT) [61] (Fig. 2A; details in Methods). We used the features obtained from each of these transformations as input to Ensemble Integration (EI), an effective framework to develop, evaluate and interpret a suite of heterogeneous ensemble-based classifiers from multi-modal data [50]. All combinations of transformations and EI classifiers were compared using a ten-fold cross-validation strategy on the High-dose (train) dataset in terms of the *F*_max_ score associated with the minority (divided) class, an appropriate metric for datasets with imbalanced classes [62, 63] and the area under the receiver operating characteristic curve (AUC) score [64, 56].

**Figure 2:**
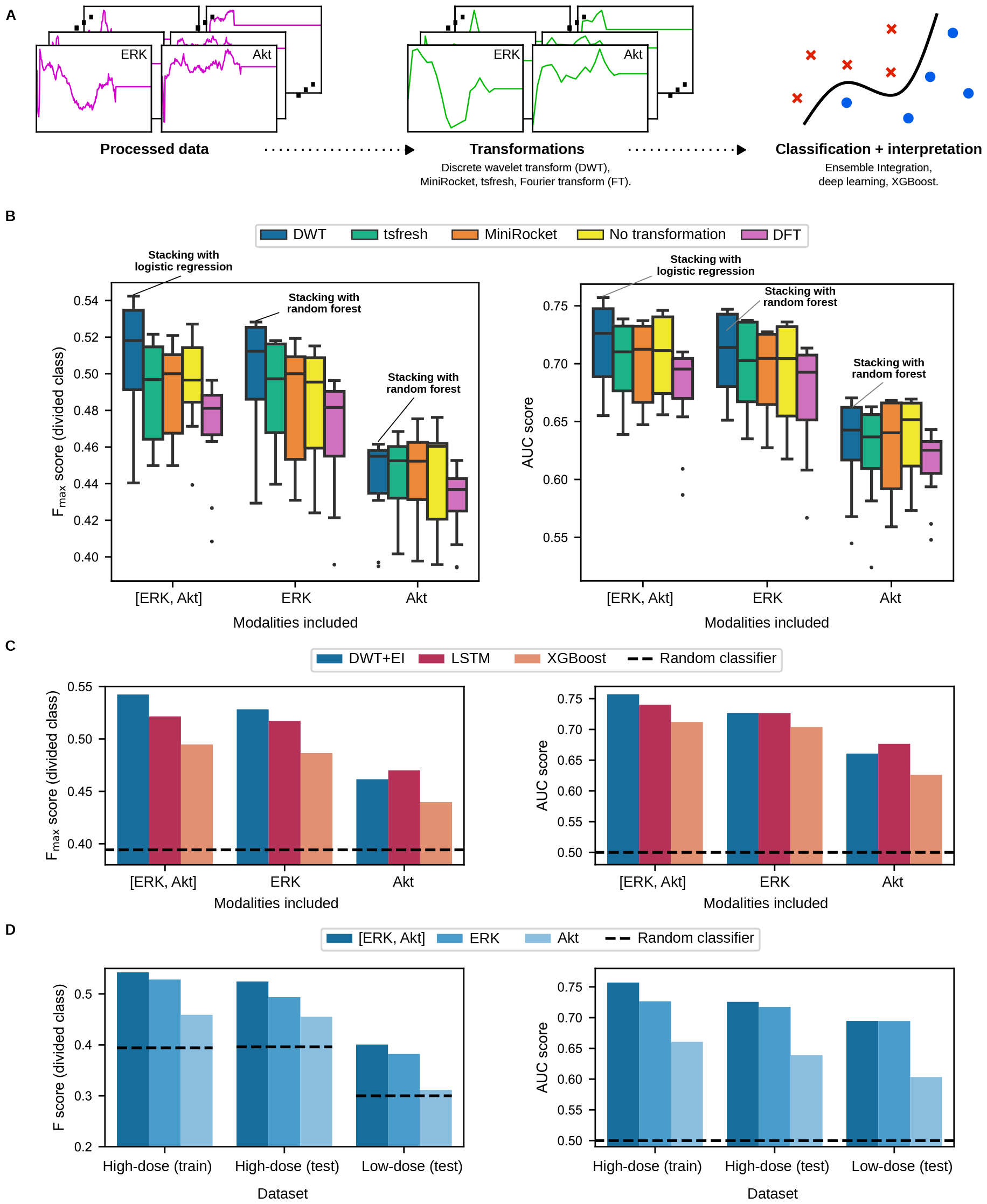
Cross-validated performance and testing of cell fate classification methods. **A**. High-level overview of the machine learning analyses used. Processed data were subjected to time course transformations to enhance the signal in the ERK and Akt time courses individually, before being used as input to several machine learning classification algorithms. **B**. Box plots showing the performance distribution of a family of ensembles developed by Ensemble Integration (EI) in combination with various transformations, as well as the multi-modal [ERK, Akt] time course. **C**. Performance of the most predictive EI algorithm combined with the Discrete Wavelet Transformation (DWT) named in **B** with arrows, as well as other established classification algorithms, namely Extreme Gradient Boosting (XGBoost) and deep learning-based Long Short Term Memory (LSTM). Both sets of results are presented in terms of the *F*_max_ and AUC evaluation measures. To calculate the *F*_max_ score, we maximized the F-measure on the training set, and then applied this threshold to discretize the test set predictions and calculate the F-measure value there. Note that the classifier with the best *F*_max_ score does not necessarily also have the best AUC score. The performance of a random classifier is shown for reference. **D**. F-measure and AUC scores of the final DWT+EI classification models on the MCF10A train and test sets. The performance of a random classifier is shown for reference. These results show that ERK is substantially more predictive than Akt across all datasets, but utilizing both ERK and Akt time courses is even more predictive.

The median *F*_max_ score of EI classifiers with the DWT was higher than other transformations for [ERK, Akt] (0.518 vs. 0.481-0.500) and ERK (0.516 vs. 0.478-0.503), and was a close runner-up for the generally less predictive Akt modality (Fig. 2B). From this, we concluded that DWT boosted the information content of the time courses the most for the prediction of cell division events using EI. We therefore selected the best DWT+EI methods, namely stacked generalization (stacking) [56] with logistic regression for [ERK, Akt] (F_max_=0.542, AUC=0.757), and stacking with random forest for ERK (F_max_=0.528, AUC=0.727) and Akt (F_max_=0.462, AUC=0.661). We used these DWT+EI combinations as the representative methods for further evaluation and model building from the respective time courses.

Next, we compared the above DWT+EI methods with other established prediction methods, namely (1) a deep learning-based long short-term memory (LSTM) network [65] and (2) XGBoost [51] on all processed time courses in the same cross-validation setup. The DWT+EI methods predicted cell fate more accurately than both the established methods in terms of both evaluation measures, especially for the more predictive [ERK, Akt] and ERK time courses (Fig. 2C). Thus, for each time course (ERK, Akt and [ERK, Akt]), we trained one final model using the corresponding representative DWT+EI method for predicting cell division on the entire High-dose (train) set, and proceeded with their evaluation on the test sets (Table 1).

Performance on the High-dose (test) set was close to that observed during training for each of the time courses in terms of both F-measure and AUC (Fig. 2D), indicating that the models were able to generalize to unseen data drawn from the same sample as the training data. For instance, the [ERK, Akt] model performed almost as well on the High-dose (test) dataset (F-measure=0.524, AUC=0.726) as it did on the training data (F_max_=0.542, AUC=0.757). The models also performed comparably on the Low-dose (test) data from a separate experiment using lower growth factor concentrations (Fig. 2D; e.g., for the [ERK, Akt] model, F-measure=0.400 and AUC=0.695). These results indicated that DWT+EI models were suitable for predicting cell division fates from ERK and Akt activity time course data in MCF10A cells.

### Combined time courses of ERK and Akt activity were the most predictive of MCF10A cell division fate, but ERK was individually substantially more predictive than Akt

Throughout our results, ERK was substantially more predictive of cell fate than Akt (Fig. 2). Furthermore, the still higher performance of the [ERK, Akt] DWT+EI model consistently showed that combined information from both time courses could improve predictive performance as compared to the individual time course classifiers (Fig. 2). This suggested complementary information among the ERK and Akt time courses, which we examined further.

### ERK and Akt activities across the time course were important for predicting MCF10A cell fate, albeit to different extents

As described above, a long-standing question in signal transduction, particularly with ERK signaling, is how signaling dynamics may relate to cell fate determination [20, 21, 22, 23, 24, 25, 26, 28, 29, 30, 31, 32, 26, 33, 23, 25, 24, 34, 22, 35, 37, 20, 38, 27, 39]. To assess if our [ERK, Akt] model could help shed light on this phenomenon, we applied an EI-associated interpretation algorithm (Supplementary Material) to identify the time points that were the most important for predicting whether a cell would divide or not. The algorithm measured importance by quantifying the change in *F*_max_ of the resultant multi-modal model when a given time point was removed from the DWT’ed sequence.

This algorithm generated an importance score for each point in the low frequency ERK and Akt time course representation, which showed that the most predictive points (high values of the importance score in Fig. 3A) were distributed throughout the time courses. These scores were generally higher for ERK (median = 0.773) than Akt (median = 0.675), consistent with the relative predictive ability of the two reporters. Furthermore, the importance scores of time points in the ERK modality were correlated with the corresponding differences in median C/N ratio between the cell fate classes, though this was not really the case for the less predictive Akt (Fig. 3B; Spearman’s rank correlation coefficient=0.449 and 0.093, respectively). These results show that ERK, and to a lesser extent, Akt activities throughout the cell cycle are associated with the likelihood of a cell division event, as opposed to directly after growth factor stimulation.

**Figure 3:**
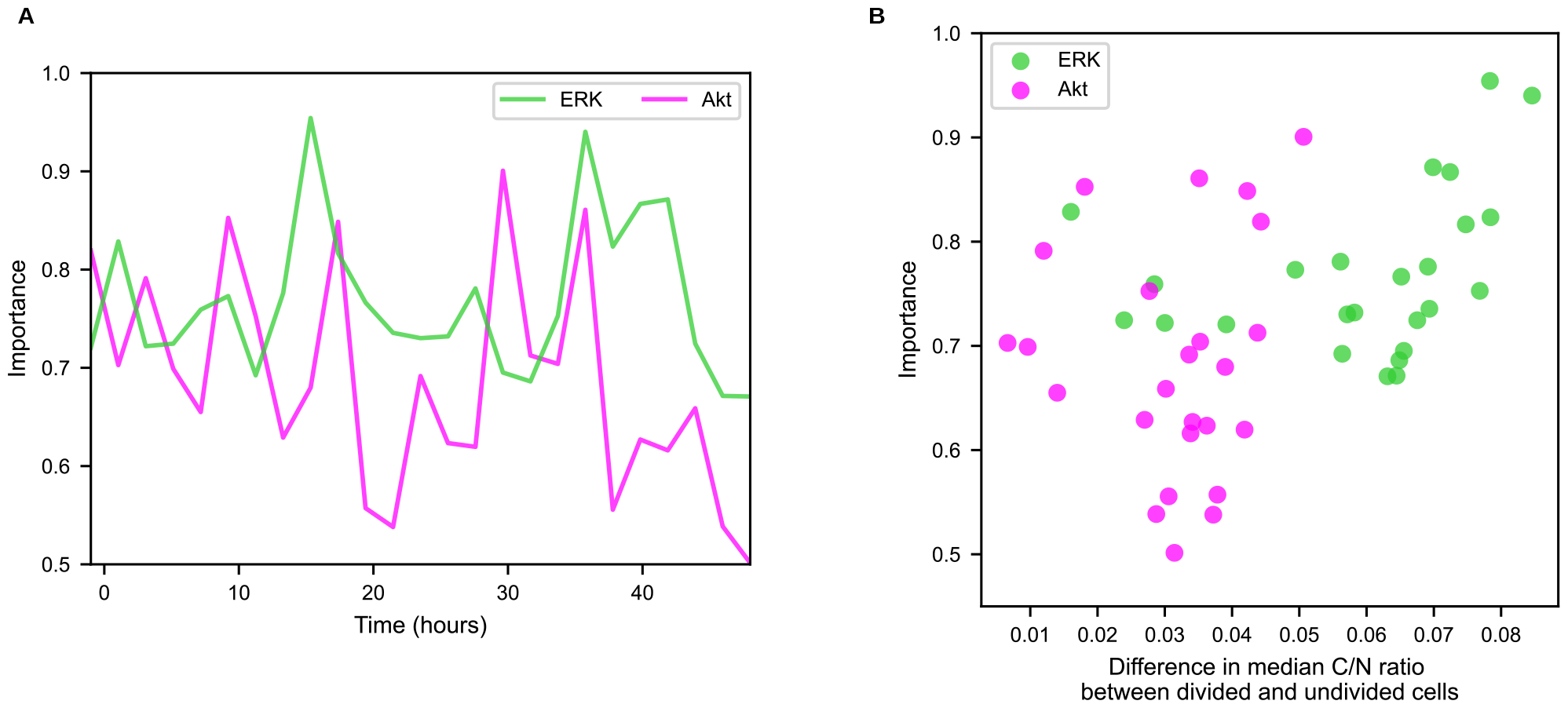
Interpretation of the multi-modal [ERK, Akt] cell fate prediction model. **A**. The respective predictive importances of ERK and Akt activities, demonstrating that time points of high importance exist throughout the time course, in particular for ERK. **B**. A scatter plot showing the importance scores of ERK and Akt activities against the differences between the median C/N ratios of the corresponding reporter for the cell fate classes (dividing and non-dividing) during the same time points. The most predictive/important time points coincided with large differences in the median C/N ratios between cell fate classes.

### The ERK model generalized to predicting cell divisions in retinal pigment epithelium (RPE) cells

Our analyses thus far focused only on MCF10A cells stimulated to divide from a serum-starved condition, and ERK activity measured using a specific KTR. This raises the question of whether the models developed from this context apply to other cell types and experimental setups. To begin to answer this question, we analyzed data from a study on retinal pigment epithelium (RPE) cells containing the EKAREN5 FRET-based ERK reporter [53]. In this study, RPE cells were not serum-starved and then treated with growth factors, but rather were asynchronously cycling in full growth medium. Applying the MCF10A-derived ERK model to this dataset (processing details in Methods) yielded F-measure and AUC scores of 0.677 and 0.694, respectively, which were comparable to those observed for the ERK time course in the High-dose (test) dataset (F-measure=0.494, AUC=0.717). Note that the higher F-measure value for the RPE (test) dataset was a result of the more balanced class distribution in this dataset, which was not the case in the other datasets we considered (Table 1). These results indicated the applicability of this model in a different biological context, as well as the utility of the DWT+EI method for model development for similar problems.

## Discussion

While the biological associations between the ERK and Akt pathways and proliferative phenotypes are established, the precise differences between the activities of these pathways in cells that do or do not proliferate are inadequately understood. In this study, we employed a data-driven approach to better understand the contributions of ERK and Akt kinase activities to growth factor-induced, single cell division fates by leveraging the natural cell-to-cell heterogeneity in these processes. To achieve this, we constructed supervised machine learning models to classify division events based on individual and combined ERK and Akt activity time courses measured by live-cell imaging of non-transformed breast epithelial MCF10A cells, a model system that is commonly used to study epithelial signaling biology and cell division control [24, 66, 67, 68, 69, 70, 3]. The results indicated that ERK and Akt activity time courses were predictive of cell division fate, especially when both the activities were considered together in the multi-modal Ensemble Integration (EI) framework [50]. Application of this model developed from MCF10A data to a different cell line (RPE), with a different experimental and biological context (asynchronously cycling cells in full growth medium), also showed that low-frequency ERK dynamics were predictive of cell division events.

We observed that processing the time courses with discrete wavelet transforms (DWTs) [58] prior to classification via heterogeneous ensembles yielded the best performance among the evaluated methods. Specifically, we applied a low-frequency DWT approximation to the time courses (Fig. S3). Our finding suggested that effective classification may not require the complete detail present in the non-transformed 15-minute interval time courses, but rather the overall trend of kinase activity described by a low-frequency representation. Moreover, an inevitable characteristic of the experimental procedure used to generate the kinase activity data used in our study was the presence of noise. Due to the ability of DWTs to effectively remove noise of different frequencies, their application proved to be particularly advantageous in mitigating its impact on classification (Fig. S3). Our results and this analysis indicate that DWTs can be a flexible, multi-resolution method to extract features from kinase activity time courses, allowing combinations of different frequency components representing variables such as cell and kinase type, as well as differences in experimental setup.

Our results also indicated that ERK dynamics were substantially more predictive of cell fate than Akt dynamics. This is consistent with the general view of the PI3K/Akt pathway being involved with growth, metabolic regulation and survival, as opposed to direct cell cycle regulation like ERK [71, 72, 73, 74, 2]. Furthermore, the multi-modal model combining ERK and Akt dynamics to predict division events was more predictive than ERK alone. This indicates that Akt dynamics had information about cell division complementary to that in ERK, which may reflect Akt’s hypothesized regulation of some aspects of initial cell cycle entry and progression [75, 76, 2], but may also suggest growth and metabolic roles.

We also found that the ERK-based model developed from the starved and growth-factor induced experimental setup was predictive of division fates from chronically growth factor treated, asynchronously cycling contexts in an independent dataset [77]. This independent dataset was from a different cell line, RPE cells, which although still of epithelial origin like MCF10A, provides further evidence of the potential generality of the models developed. Recent live-cell imaging studies of ERK signaling in epithelial cells have instead looked at confluent cells in chronic growth factor treatment conditions, arguing that such a setup is closer to the epithelial biology that would be observed in tissues [24, 23]. Large annotated datasets linking ERK dynamics to cell division events in confluent, chronic growth factor contexts would be needed to understand how different such scenarios truly are. While obtaining the imaging data themselves is relatively straightforward, high-quality annotated datasets like ours still often require substantial manual effort. Advances in machine learning for image analysis may be able to help generate such datasets more efficiently [78, 79, 80].

Another question we were able to address using the developed models was which time points of signaling dynamics were the most predictive of cell division events. As discussed above, there are multiple hypotheses for such relationships, including transient versus sustained signaling, the frequency and/or magnitudes of pulsing and time-integrated activities [20, 25, 22, 26, 34, 38, 27, 37]. In the data analyzed here, there was not substantial evidence of pulsing, but this does not mean it might not be predictive in other contexts. Based on a systematic model interpretation algorithm, our results suggested that no single time point in the ERK and Akt time courses was solely more useful than others for predicting cell division events. This is more consistent with a time-integrator model, whereby the amount of time the pathway is on dictates the probability of cell division [37, 20, 38, 27, 39, 27]. Recent studies in MCF10A cells have suggested such a relationship between ERK signaling and cell cycle progression [38]. However, proving such a relationship to be causal is challenging, since cell cycle progression depends on the activity of the ERK pathway. Optogenetic approaches are a suitable tool for such examinations, and have been used to establish relationships between ERK signaling and cell fates in other contexts [3, 25, 20, 35].

There are multiple considerations for experimental design that could have a bearing on interpretation. Foremost, while the trained models were shown to be predictive, they were not 100% accurate, implying that other biological sources of noise, such as p53 / DNA damage [40], play operational roles in predicting cell division fate. In our MCF10A cell studies of Akt dynamics, large, saturating doses of insulin were used, because that is the standard for MCF10A growth media. Insulin is a strong activator of the Akt pathway [81, 82, 83]. If Akt activity was consequently very high, its fluctuations over time in a single cell may not dip below thresholds that interfere with its ability to drive cell cycle progression. Thus, growth factor conditions that do not as strongly activate the Akt pathway may lead to a different conclusion about the relative importance of ERK activity versus that of Akt for driving cell division events. Yet, we did also test the predictive models on a dataset with 10-fold lower growth factor doses, but the conclusions were similar. This could mean that insulin concentrations remained high, but it could suggest that the original interpretation of Akt activity dynamics being less predictive are more likely to be true. There are several growth factors that induce both cell division and ERK and Akt activities. Whether the predictive relationships between ERK and/or Akt dynamics and cell division change depending on what activates the pathways remains an open question. Besides growth factor dose and type, there are other important experimental elements to consider in the context of our results. In the MCF10A experiments, somewhat sparsely seeded cells were serum- and growth factor-starved prior to the experiment, treated with growth factors, and then observed for 48 hours. Observing cells for longer could have revealed more division events, but also confounded results, as cells dividing early in the time course could have gone through the same process again. Thus, our results may be more strictly interpreted as most effective for predicting the fates of cells most likely to divide (relatively) quickly. While serum and growth factor starvation is a long-established mode of studying cell signaling [84], chronic treatment with growth factors may create a different relationship between signaling dynamics and cell division fates [4, 24, 12]. Yet, the models developed from starvation experiments in MCF10A cells were found to be predictive for RPE cells asynchronously cycling in full growth media, providing at least one example of generality.

In summary, our results suggest that discrete wavelet transforms of cell signaling dynamics data, combined with ensemble-based classification models, may be a generally useful tool for cell fate prediction. Our particular application to cell division associated with ERK and Akt dynamics suggested some generality of the relationship between these dynamics and cell division for epithelial cell lines in both acute and chronic growth factor conditions. Model interpretation suggested that the time-integrator model of how these pathways influence cell division fate is more likely in the studied contexts. The availability of more annotated datasets would enable a more expansive study to understand how general such relationships are in different cell types, growth factors, doses and other experimental setups.

## Methods

### Dataset sources and availability

Our analysis and conclusions are all reproducible by following the instructions in the GitHub repository: https://github.com/GauravPandeyLab/predicting-cell-division.git. MCF10A datasets used in this study were the dual-reporter datasets presented in a previous paper [27], and are freely available at the repository mentioned therein. The RPE datasets were produced independently [53], and were obtained by contacting the authors.

### Dataset processing

#### MCF10A datasets

The MCF10A datasets were generated as described in detail previously [27]. Briefly, cells expressing reporters were seeded, allowed to attach overnight, and serum- and growth factor-starved for 24 hours. An hour of baseline images were taken before applying growth factor treatment, after which, images were taken every 15 minutes over the next 48 hours, giving a total of 49 hours of measurements. The resultant image time courses were analyzed to measure kinase activity (cytoplasmic to nuclear fluorescence ratio of the kinase translocation reporter (KTR) (C/N ratio)). Experiments in this study included biological replicates for reproducibility.

To prepare sufficiently sized sets of examples of labeled cells to train and test machine learning methods/models, we merged the data across replicate experiments from the original study after confirming that there were no batch effects (Fig. S1). In the resultant data, experiments with a high-dose of growth factors induced more divisions, resulting in a less severe cell fate class imbalance than that in low-dose settings (24.6% divided cells for high-dose data compared to 16.6% for low-dose). To have sufficient representation of the divided class in the data, we used the high-dose data only for both the assessment of candidate methods for cell fate classification and the subsequent final model training. We then performed an 80:20 train/test split on the high-dose data, yielding one training and two test MCF10A datasets (Table 1).

Processing steps (see Supplementary Material) were performed on the training set, and any processing parameters derived were then applied during processing of the test sets.

#### RPE dataset

The RPE experiments (part of a larger study [53]), in addition to using a different cell line, only measured ERK activity with the EKAREN5 reporter (and not Akt, or with a KTR reporter), had cells asynchronously cycling in full growth media (not initially serum-starved), and included measurements over a 4 day period with a 10 minute frequency and multiple division events along the time course (versus 2 days, 15 minute frequency, and the first division event in the MCF10A datasets). Key differences are summarized in Table 1. This study contained multiple experimental setups with different combinations of treatments. The RPE (test) dataset used in our study was generated from the experiment with no DOX and 62.5nM ERK inhibitor (ERKi), because this was the only condition where cell division fate heterogeneity was observed in the last 49 hours of the time course.

To be able to test the ERK model developed from the MCF10A dataset in this different setup, we labeled the RPE cells as divided/undivided according to whether a division occurred in the last 49 hours of their respective time courses. This also allowed us to remove cells whose division events early in their time courses reflected those committed to division prior to treatment with ERKi. We linearly interpolated each time course in RPE (test) to match the 15 minute measurements of the MCF10A datasets. We then applied the same processing steps as those described above for the MCF10A datasets. To account for different reporters, we scaled ERK activity in the resultant RPE (test) dataset to make it as consistent as possible with the measurements in the High-dose (train) dataset (details in Supplementary Material).

### Classification methods and their evaluation

We evaluated a number of classification methods using a stratified, ten-fold cross-validation procedure [85] applied to the High-dose (train) dataset. We addressed class imbalance in this process by undersampling the majority (undivided) class during training, and evaluated the performance of the tested methods using the *F*_max_ score associated with the minority (divided) class [62, 63]. This measure is defined as the maximum value of the F-measure across all classification score thresholds, allowing each classification method to achieve their most effective performance. We also report the area under the receiver operating characteristic curve (AUC) score [64, 56].

We compared several approaches for classifying cell division fate in the above evaluation setup (details of the approaches in Supplementary Materials). The first approach was to extract features using a time course transformation (next subsection) prior to building predictive ensembles using the Ensemble Integration framework [50], which is designed to flexibly and effectively integrate multi-modal data like the ERK and Akt time courses. Secondly, we considered a deep learning-based Long Short Term Memory (LSTM) [86] classification method whose architecture was specifically developed to handle sequential dependencies in the time courses. Finally, we considered eXtreme Gradient Boosting (XGBoost; [51]), which has been found to be the most effective performer in many classification tasks [87, 88, 89].

A particular interest of our study was to compare multi-modal methods/models that utilized information from both ERK and Akt time courses to those built only using the information in the individual time courses. Thus, we evaluated all the classification methods considered on all three modalities, i.e, ERK, Akt and [ERK, Akt], with the aim of building three different final models. For each of the modalities, we calculated the median F_max_ scores of the EI methods tested for each transformation. In addition to analyzing the performance trends across the transformation+EI method combinations, we also selected the best-performing combination for each modality for final model building. The LSTM-based neural network and XGBoost performances were similarly evaluated and compared.

### Time course transformations

To more effectively extract information from the ERK and Akt time courses prior to cell fate classification, we subjected them to a selection of transformations [57] before evaluating their predictive capabilities using EI. We used Minirocket [59] to generate features of random convolutions, since this method has been highly successful across a range of applications [57]. Statistical feature extraction is also a common method for classifying time courses [90, 91] and so we applied the tsfresh [60] package to automatically generate a large number of statistical features characterizing the time courses. We also evaluated more classical transformations in the form of the Fourier transform amplitude (see for example, [61]) to gauge the magnitude of different frequency components present in the time courses. Finally, we used discrete wavelet transforms (DWTs, [58]), which, unlike Fourier transforms, can facilitate feature extraction with both time and frequency localization.

### Ensemble Integration

We utilized Ensemble Integration (EI) [50, 92] to compare several interpretable ensemble methods and train our final models based on the best performers. Within the EI process for each modality, we trained a selection of base classifiers, and combined their predictions via an ensemble classifier. The specific ensemble classifier was selected from a variety of ensemble algorithms via a nested cross validation. Although EI is able to build ensembles from single data modalities (i.e. ERK or Akt alone), it is especially effective for data from multiple sources (multi-modal data), the [ERK, Akt] modality in this study. The exact base and ensemble predictor algorithms and parameters that were used in this study can be found in Supplementary Material.

### Interpretation algorithm for EI models

We calculated an importance score for the time points of the discrete wavelet-transformed time courses constituting the best performing [ERK, Akt] model (Fig. 3), i.e., stacked generalization with logistic regression (Results). The algorithm used for this calculation utilized a randomized permutation procedure [93] to calculate feature ranks of each base classifier algorithm, before applying a weighted average over all the time points based on the learned weights of the logistic regression stacker (Algorithm S1). The output of this algorithm was an importance measure between 0 and 1 for each time point, where values closer to 1 indicate higher importance (more details in Supplementary Material).

## Acknowledgements

We thank Jia-Yun Chen, Peter Sorger and Galit Lahav for sharing the RPE dataset generated in their study, as well as associated information. We also thank Alexander Davies and Rawan Makkawi for helpful discussions.

