## Supplementary material for "Low-frequency ERK and Akt activity dynamics are predictive of stochastic cell division events"

### Batch effects among replicate datasets

All MCF10A datasets used in our study had replicates, which could be combined to increase the examples available for training and testing our machine learning methods/models. However, before doing this it was important to determine if there were batch effects between those replicate dataset. We conducted a principal component analysis (PCA) of the combined replicate data associated with high and low dose experiments, and found no evidence of batch effects (Fig. S1), and so, we merged the data from the replicates.

### Processing of the datasets

One consequence of the experimental procedures that generated the MCF10A datasets was that kinase activity measurements ceased when a cell divided. This meant that undivided time courses contained a fixed number of measurements over the 49 hour duration of the experiment, whereas divided courses contained less. ML algorithms often require fixed length input time courses. Padding, i.e., adding zeros or another pre-specified value at the end of shorter time courses, is a commonly used approach to achieve this. However, in our case, the existence of padding alone would have been indicative of the divided class.

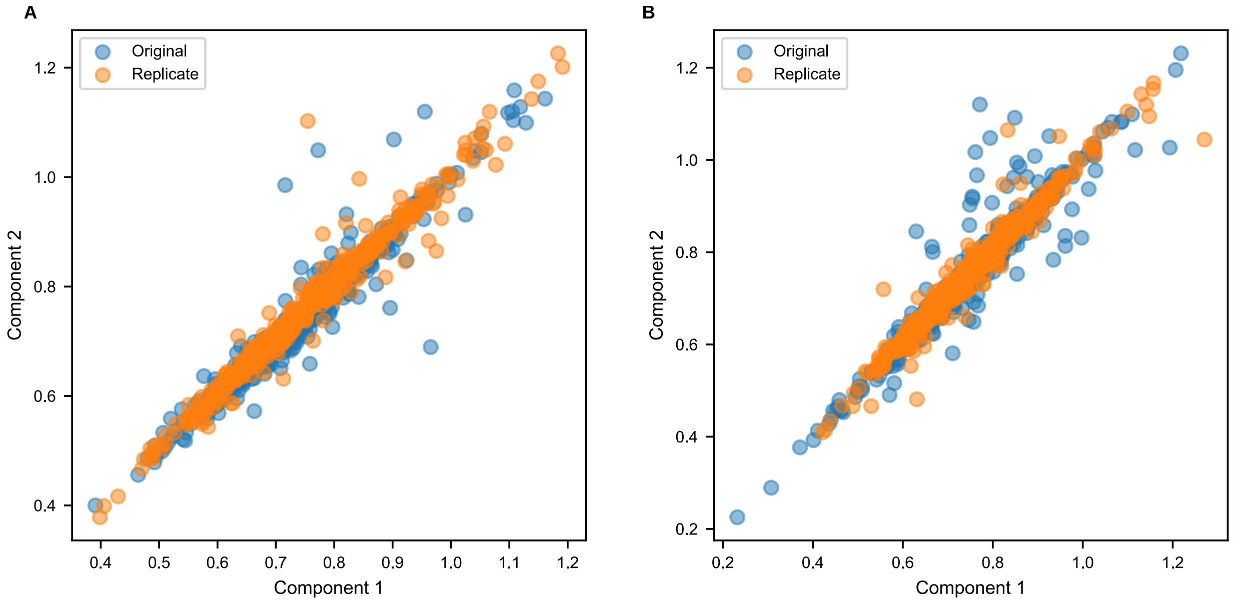


**Figure S1:** Assessment of batch effects among replicate datasets using Principal Components Analysis (PCA). Similar clustering in the principal components implied that there are minimal batch effects between original and replicate experiments for A. High-dose data and B. Low-dose data. We therefore combined original and replicate data to increase the number of examples in this study and enhance the robustness of the findings.

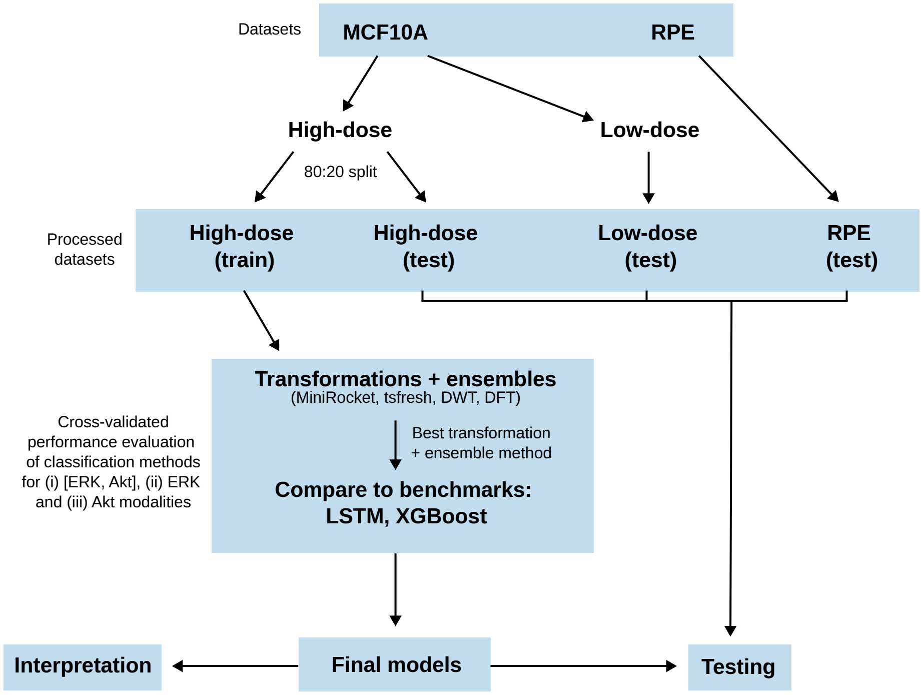


**Figure S2:** Workflow overview of the analysis performed in this study.

We therefore truncated undivided time courses to match the length distribution of the divided ones in the training set before applying padding. This was done by randomly sampling from the discrete distribution of lengths of divided cell time courses in the High-dose (train) dataset, and randomly assigning them to the time courses from the undivided cells. For each undivided time course, we removed all time points beyond this sampled length, and then assigned the mean of the time course values until this point (mean padding) to its right hand side so that all time courses–divided and undivided–were of equal length. Note that the ERK and Akt time courses corresponding to a single divided cell were of the same lengths; we therefore sampled lengths from one modality and applied it to both modalities of an undivided time course.

The RPE (test) dataset had several differences in the experimental process from the MCF10A data, including a different reporter (Table 1). We therefore scaled the values in RPE (test) to be consistent with those in High-dose (train). We did this by finding the median values in the time courses in both datasets to reduce the impact of outlier values, and calculated the midpoint of the range, i.e. $[\max(\text{median}) + \min(\text{median})]/2$. We then linearly scaled the time courses in the RPE (test) dataset so that this resultant midpoint matched that of High-dose (train). Implementation details are available in the code (`https://github.com/GauravPandeyLab/predicting-cell-division.git`).

### Cross-validated performance analysis and model selection

We partitioned the High-dose (train) dataset into 10 folds that were kept consistent in the evaluation of all methods tested. To reduce the effects of class imbalance, we undersampled all training folds to equalize the sizes of the divided and non-divided classes. Our analysis considered three separate cases, using (i) only ERK time courses, (ii) only Akt time courses, and (iii) both ERK and Akt time courses, with the aim of building three final models to evaluate on our test datasets. Classification algorithms were trained on each set of nine training folds, and used to make predictions on the remaining test fold. Predictions across all test folds were concatenated before being evaluated. Evaluation of classification performance was based on the $F_{\max}$ score associated with the minority divided class. This measure, which is the maximum value of the F-measure across all classification thresholds, has been suggested to be more reliable than other more commonly used evaluation metrics for unbalanced class scenarios like ours [1, 2]. We also calculated the area under the receiver operating curve, and for reference, the performance of a random classifier in our results (Fig. 2B-D).

At the beginning of the above evaluation process, a number of transformations were evaluated by assessing the predictive performance of classifiers built using Ensemble Integration (EI) [3]. Transformations were applied to ERK and Akt modalities separately to generate features which were used as input to EI, which

| Method | Base | Ensemble |
|---|---|---|
| Logistic regression [6] | ✓ | ✓ |
| Gaussian naive Bayes [6] | ✓ | ✓ |
| k-nearest neighbours [6] | ✓ | ✓ |
| Support vector machine [6] | ✓ | ✓ |
| Decision tree [6] | ✓ | ✓ |
| Random forest [6] | ✓ | ✓ |
| Gradient boosting [7] | ✓ | ✓ |
| Extreme gradient boosting [5] | ✓ | ✓ |
| Multi-layer perceptron [6] | ✓ | ✓ |
| Adaptive boosting [8] | ✓ | ✓ |
| Ensemble selection* [9] | ✗ | ✓ |
| Mean* | ✗ | ✓ |

**Table S1:** Component methods of ensembles applied using Ensemble Integration [3]. Each ensemble generated utilized all base classifiers listed. Unless starred the ensemble method is applied using stacked generalization [6]. Specific parameters/hyperparameters used are given in the GitHub repository.

carried out the above cross-validation process internally, and generated performance scores for each ensemble method considered, forming a distribution of performance scores (Fig. 2B). The most effective transformation was identified as the one with the highest median EI performance across cases (i)-(iii), and selecting the most frequent one. The final transformation + ensemble method for each course was determined to be the one with the highest $F_{max}$.

This best-performing method was compared to two benchmark algorithms: an LSTM-based [4] deep learning algorithm, and XGBoost [5]. In the case of the LSTM model, we further partitioned each (outer) training fold into 10 inner folds (nested cross validation) to optimize the number of training epochs and avoid overfitting (a common issue with neural networks especially with a small number of samples). For each outer fold, inner test folds were used to track the validation loss at each successive epoch, and the epoch with the lowest loss was stored. The median of these epoch values was then taken across the inner folds. An LSTM model was then trained (with this median epoch value) and evaluated on each time course and their combination in the same cross-validation setup as described above.

For the best performing methods for cases (i)-(iii), we trained final models on the full High-dose (train) dataset (all ten folds). We then evaluated these models on the held out test sets (Table 1). A more detailed overview of the analysis performed in this study is shown in Fig. S2.

### Ensemble Integration configuration

We used the recent Python eipy package [10] to train and evaluate ensembles using the EI framework [3]. Table S1 lists the base and ensemble classification algorithms used within EI in this study.

### Discrete wavelet transforms

Discrete wavelet transforms (DWTs) are a principled approach to decomposing time series into time and frequency components simultaneously [11], unlike discrete Fourier transforms which represent time series in the frequency domain alone. The DWT decomposes the signal into "approximation" and "detail" coefficients. The approximation coefficients represent the coarse, low-frequency components of the signal, while the detail coefficients capture the high-frequency details.

We took the approach of extracting low frequency information from the signal by performing repeated DWTs as a cascade (Fig. S3). We applied three Haar DWTs to each processed time series example successively, discarding the high frequency detail component of the signal at each stage. The resulting low frequency representation had time points spaced approximately 2 hours apart.

### Interpretation of cell fate prediction models

We interpreted the final [ERK, Akt] model as follows. We began by applying a permutation feature importance algorithm [12] to each base classifier model across both modalities. This algorithm (see Algorithm S1) evaluates the change in $F_{max}$ score when randomly shuffling a feature column for a particular time point, thereby

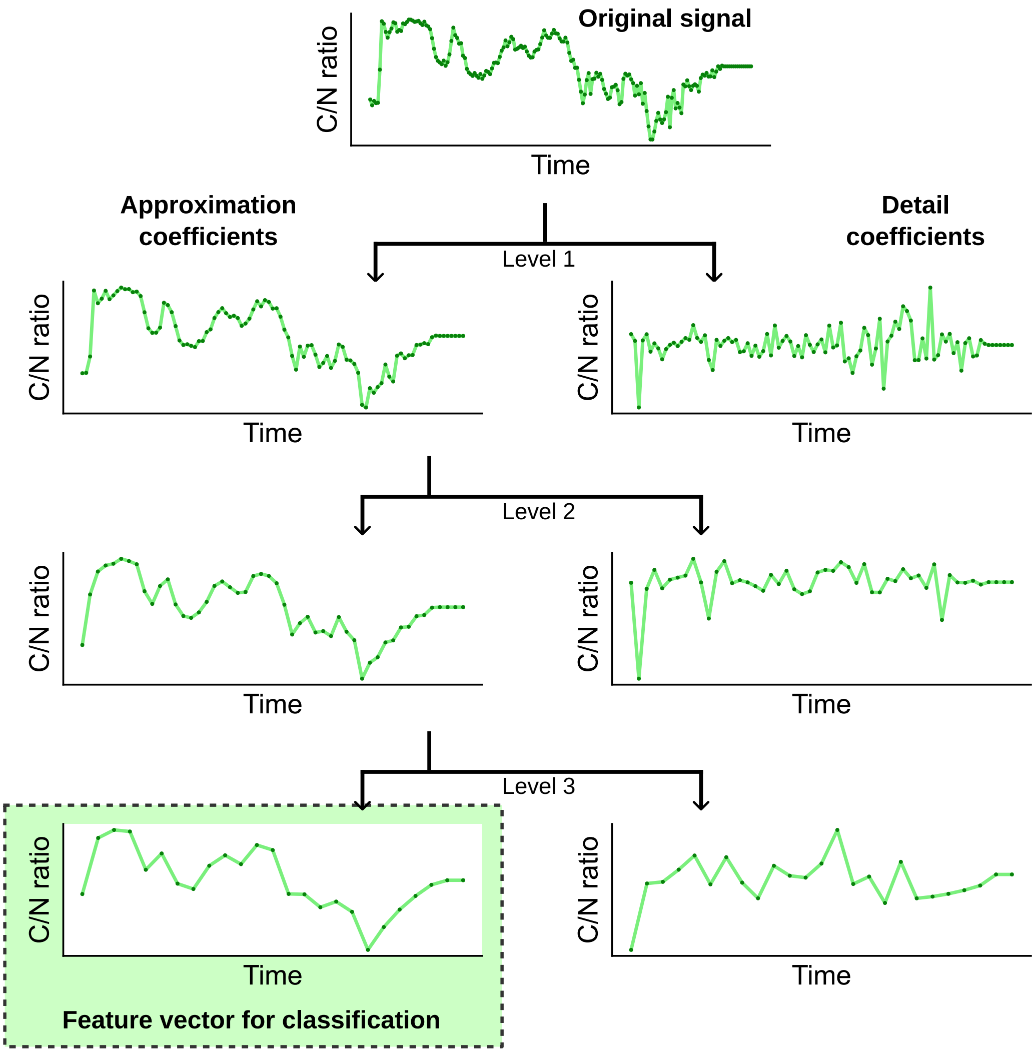


**Figure S3:** Feature extraction from time courses using the discrete wavelet transform (DWT). We applied a Haar DWT to the original signal to produce the first level of approximation and detail coefficients. We then applied successive transformations to the approximation coefficients two more times as a cascade to obtain level 2 and 3 approximation and detail coefficients. We selected the level 3 approximation coefficients as features for input to our classification models, since these coefficients retained the low frequency shape of the signal, while removing higher frequency information including noise. This representation of the signal contained only 25 data points as opposed to 198 in the original signal, thus also helping with dimensionality reduction.

breaking the relationship between the measurement at that time point and the target class, yielding a measure of the former's overall importance for making predictions. This process was repeated several times for each time point, and the mean difference in performance recorded. These importance means were then percentile ranked. Thus, we obtained a set of local feature (time point) ranks (LFR) for each base classifier in the ensemble.

The LFR were independent of other base classifiers and, therefore, modalities. We needed to weight and combine the LFR according to the importance of the base model predictions in order to calculate the importance score of each time point for the final ensemble model. This model was based on the DWT+EI logistic regression stacker (Results). Since the ensemble algorithms took probabilities as input (i.e. all features had the same scale), we were able to gauge the importance of the base classifiers in the EI model by assessing the magnitude of the learned coefficients of the logistic regression stacker. Thus, we percentile ranked the absolute values of logistic regression coefficients, and used them as weights in a weighted average applied to the LFR across base classifiers. This resulted in an importance score for each time point which varied between 0 and 1, with higher scores indicating greater importance.

**Algorithm S1** Computation of importance scores for the [ERK, Akt] ensemble model. This model used stacked generalization with logistic regression. The algorithm utilizes `sklearn.inspection.permutation_importance` from the scikit-learn machine learning library [14], which is a model agnostic interpretation method.

**Data**:
`n_base_predictors`: number of base predictors in the ensemble.
`n_features`: number of features across all modalities.
`base_predictors`: list of base predictors in the ensemble.
`lr_coeffs`: vector of length `n_base_predictors` containing the learned coefficients in the logistic regression stacker. The larger the magnitude of the coefficient, the more important the corresponding base predictor.
**Functions**:
`permutation_importance`: the function `sklearn.inspection.permutation_importance`, which outputs an feature importance measures for a single model.
**Result**:
`weighted_importance`: vector of length `n_features` containing feature importance scores between 0 and 1, weighted according to the contribution of base predictors in the ensemble.

```
Calculate local model ranks (LMR) and local feature ranks (LFR):
LMR ← vector of length n_base_predictors, containing percentile ranks of abs(lr_coeffs).
LFR ← array of shape (n_features × n_base_predictors).
for m ← 1 to n_base_predictors do
    LFR[:,m] ← percentile rank permutation_importance(base_predictor[m])
Calculate weighted_importance:
weighted_importance ← empty vector with length n_features
for i ← 1 to n_features do
    for j ← 1 to n_base_predictors do
        rank_product_importance ← LMR[j] × LFR[i,j]
    weighted_importance[i] ← mean(rank_product_importance[i,:])
```

### LSTM benchmark

There is substantial flexibility when constructing LSTM architectures, but we found that configurations with large numbers of parameters were prone to overfitting, which was likely due to the relatively small amount of training data available for this task. For this reason, we limited the number of layers in our LSTM benchmark, as well as the number of units in each layer. This benchmark consisted of two hidden layers: a bi-directional LSTM layer, followed by a fully connected layer with ReLU activation. Each of these layers contained 64 units. We also used heavy dropout to regularize the network to further control overfitting.

We optimized the binary cross-entropy loss used in the benchmark using Adam optimization [13]. Full details of the method used can be found in the GitHub repository.
